# Platycodin D2 Targets Vimentin to Suppress Fibroblast-Myofibroblast Transition: A Novel Therapeutic Strategy Against Pulmonary Fibrosis

**DOI:** 10.64898/2026.11.03.763586

**Authors:** Xianshengjie Lang, Zheng Ma, Chuning Yao, Xinyu Miao, Yuchenyang Kao, Wenwen Wang, Shuangli Sun

## Abstract

**Background:** Pulmonary fibrosis is a progressive lung disease characterized by excessive extracellu-lar matrix deposition and irreversible decline in lung function with limited effective therapeutic targets. Vimentin, a type III intermediate filament protein, plays a critical role in fibroblast activation and fibrogenesis, representing a promising drug target.

**Methods:** Novel vimentin ligands were screened via virtual screening. SPR assay verified compound-vimentin binding affinity. CETSA and CD spectroscopy analyzed protein structural changes after compound binding. In vitro FMT model and in vivo bleomycin-induced pulmonary fibrosis model were used to assess antifibrotic effects, and mutagenesis assays identified key binding amino acid residues.

**Results:** Platycodin D2 (PD2) from Platycodon grandiflorus was identified as a novel high-affinity vimentin ligand (Kd = 392.8 nM). PD2 binds vimentin directly, alters its secondary structure and induces insoluble aggregation. It effectively inhibits TGF-β1-induced FMT in vitro and mitigates bleomycin-induced pulmonary fibrosis in vivo, reducing collagen deposition and improving animal survival. Critically, mutagenesis val-idation confirmed that key residues (Glu138 in chain A, Glu132 and Glu143 in chain B, as well as Glu50 and Arg54 in chain C) are essential for this interaction.

**Conclusions:** Our findings establish PD2 as a novel pharmacological inducer of aberrant vimentin aggregation, presenting a distinct target-driven strategy for antifibrotic drug development.

## Introduction

Pulmonary fibrosis (PF) is a chronic, progressive lung disease characterized by irreversible scarring and excessive deposition of extracellular matrix, leading to impaired gas exchange and progressive respiratory failure^1^. PF carries a rising global burden, and the COVID-19 pandemic has further elevated its incidence and severity; 20–30% of severe COVID-19 patients develop irreversible lung fibrosis, greatly exceeding the complication rate of earlier viral pneumonias^2,3^. These findings highlight an urgent and growing need for effective therapeutic interventions for PF. Despite this growing need, the therapeutic armamentarium for PF remains limited. Although pirfenidone and nintedanib are approved by the FDA for the treatment of PF, long-term use of these agents is associated with adverse events, such as allergic reactions and arterial thromboembolism^4^. Currently, 28 small-molecule anti-PF compounds are under clinical investigation globally, but only three have progressed to Phase III registration trials, underscoring the formidable challenges and slow trajectory of drug development in this area^5,6^. Consequently, there remains a critical unmet need for novel, disease-modifying therapies against PF.

Vimentin, a member of the type III intermediate filament family, is widely expressed in mesenchymal-derived cells, including fibroblasts, endothelial cells, and immune cells^7^. Beyond its canonical role in maintaining cytoskeletal integrity, vimentin functions as a central regulator of cell migration, mechanosensation, and signal transduction^8^. Critically, vimentin is a hallmark marker of epithelial–mesenchymal transition, where it drives the expression of fibrosis-related proteins and promotes extracellular matrix deposition^9^. The biological functions of vimentin depend on a dynamic cycle of assembly and disassembly. Soluble vimentin tetramers serve as the functional unit, polymerize to form insoluble intermediate filaments, which are essential for vimentin-mediated cytoskeletal organization^10–12^. Under physiological steady-state conditions, the mature insoluble vimentin filaments can be depolymerized into soluble tetramers, which participate in signal transduction^13–15^.

Given its pivotal roles in fibrosis, vimentin has attracted considerable interest as a potential therapeutic target^16^. The natural product Withaferin-A (WFA) has been shown to disrupt vimentin filament networks, leading to suppression of tumor metastasis and attenuation of organ fibrosis^17^. Studies have indicated that targeting vimentin can alleviate PF^18–20^; however, the clinical translation of WFA has been hindered by its broad cytotoxicity and narrow therapeutic window. These limitations underscore the need for alternative vimentin-targeting strategies with improved selectivity and safety profiles. Although targeting vimentin has shown promise in alleviating pulmonary fibrosis in experimental models, no vimentin-based therapies have entered clinical trials, and the underlying molecular mechanisms remain incompletely understood.

To develop effective novel vimentin-targeting compounds with therapeutic potential for pulmonary fibrosis, we performed virtual screening of a library of traditional Chinese medicine (TCM) compounds using vimentin as the molecular target. This screen identified Platycodin D2 (PD2) as a potential ligand of vimentin. PD2, an oleanane-type triterpenoid saponin and one of the major bioactive constituents of *Platycodon grandiflorus* root, has been reported to inhibit the TGF-β/Smad and NF-κB signaling pathways^21,22^. It exhibits well-documented preclinical activities, including expectorant, anti-tussive, anti-pulmonary fibrotic, hepatoprotective, antitumor, anti-inflammatory, and immunomodulatory effects^23^. Despite these promising pharmacological profiles, the direct molecular target of PD2 and its mechanism of action in fibrosis have not been defined.

In this study, we demonstrated that PD2 binds to vimentin and induces a conformational change that disrupts its normal assembly dynamics, driving the protein from the soluble state into an aberrantly aggregated insoluble state. This aggregation-dependent depletion of functional vimentin blocks fibroblast-to-myofibroblast transition (FMT), reduces collagen deposition, and ultimately ameliorates bleomycin-induced pulmonary fibrosis in vivo. Our findings establish PD2 as a pharmacological inducer of vimentin aggregation and reveal a previously unrecognized mechanism—ligand-induced aberrant aggregation of a pro-fibrotic effector—that may offer a new strategy for antifibrotic drug development.

## Materials and Methods

### Cell culture

The lung adenocarcinoma cell line A549 (generously provided by Dr. Jia Rui, Shandong University) was maintained in DMEM/F12 (PM150312, Procell, Wuhan, China) supplemented with 10% FBS (FSP-500, ExCell, Shanghai, China). The human lung fibroblast MRC-5 cell line (MRC-5) were purchased from the cell bank of the Chinese Academy of Sciences (Shanghai, China). Cells were cultured in DMEM (C11995500BT, Gibco) supplemented with 10% FBS. All cells were cultured at 37 °C in 5% CO2 and 95% humidity environment.

### Cell viability

A549 cells were seeded in 96-well plates at a density of 2 x 10^3^ cells per well. After 24 hours of incubation, PD2 was dissolved in DMSO and diluted stepwise in serum-free medium to the target concentration (0.1-100 μM, final DMSO concentration <0.1%). The original medium was discarded, and the experimental group was treated with drug-containing medium, the control group was added with an equal volume of DMSO dilution, and the blank group was added with medium only. At least three wells were set up in each group, and the intervention was carried out for 24, 48, and 72 hours, respectively. At the end of the intervention, 10 μl of CCK-8 reagent (C0041, Beyotime) was added to each well, and incubated for 2 hours away from light. The absorbance (OD) was measured at 450 nm using a microplate reader (BioTek SynergyH1). The experiment was repeated independently three times.

### Cell wound healing assay

MRC-5 cells were seeded into 6-well plates at a density of 1.0-1.6×10^5^ cells/well and cultured until 90-95% confluence. A sterile 200 μl pipette tip was used to scratch the monolayer perpendicular to the well plate, and the detached cells were gently washed away with PBS and replaced with serum-free basal medium DMEM (without or with 5 ng/ml of TGF-β) and different concentrations of PD2(0, 5 and 10 μM). Fixed-position scratch images were taken under an inverted phase contrast microscope (×100, calibrated scale) at 0 and 48 h, respectively.

### Cell immunofluorescence assay

Glass coverslips were placed in 12-well plates, and MRC5 or A549 cells were cultured for 24 hours. TGF-β was added to induce fibrosis. Following this, 5 μM PD2 or control DMSO (vehicle) was administered for a 48-hour intervention. Cells were fixed in 4% paraformaldehyde for 10 min, permeabilized with 0.1% Triton X-100 (CAS 9002-93-1; Sangon Biotech, Shanghai, China) for 10 min, and blocked with 2% bovine serum albumin (BSA; ST023-50g, Beyotime Biotechnology) for 30 min. After two washes with PBS, cells were incubated overnight at 4°C with primary antibodies against Fibronectin, Collagen I, α-SMA, or vimentin (1:100 dilution). Subsequently, cells were washed three times with PBS (5 min per wash) and incubated with species-specific fluorescent secondary antibodies (1:500 dilution) for 1 h at room temperature in the dark. Following three additional PBS washes (5 min each), excess liquid was gently blotted from the slides using filter paper. A drop of anti-fade mounting medium containing DAPI was applied, and coverslips were carefully placed over the slides to avoid air bubbles. Fluorescence images were acquired using an FV3000 laser scanning confocal microscope (MEAGLE 320-PRO; Changyi Optics).

### Animals

C57BL/6 mice were purchased from Vital River Laboratory Animal Technology, Co., Ltd. (Beijing, China). Mice were housed under controlled conditions (temperature 22-26°C, humidity 40-70%) with a 12-hour dark/light cycle. All animal care and experimental protocols were approved by the Ethics and Use of Animal Care Committee of Shandong University and were conducted according to the guidelines of the committee.

### BLM-Induced Pulmonary Fibrosis in Mice

Thirty 6-8-week-old male C57BL/6 mice were randomly divided into three groups (n = 10 per group), including the control group (the control group was also the sham surgery group), the bleomycin sulfate-induced group (the BLM group), and the bleomycin sulfate-induced, 5 mg/kg PD2-intervention group (the BLM+PD2 group). Mice were anesthetized by inhalation of isoflurane, and then the trachea and esophagus were separated with a blunt instrument. To induce PF, mice were given a single intratracheal injection of BLM (5 mg/kg body weight in 0.05 ml of sterile saline; B802467, Marcolin, China). Meanwhile, mice in the sham group received an equal volume of sterile saline in the same manner. Mice in the BLM+PD2 group were injected intraperitoneally with 5 mg/kg of PD2 once a week, and other mice were injected intraperitoneally with an equal volume of saline as a control. On the 21st day after BLM injection, mice in each group were anesthetized and lung tissue was rapidly excised after sacrifice. A portion of the right lung was fixed in 4% paraformaldehyde, and a portion of the left lung tissue was stored at −80°C for subsequent real-time-PCR and Western blot analysis. The body weights of the mice were recorded daily.

### Histological Examination

After mouse euthanasia, the left lung was removed and the tissue was washed clean with PBS and fixed in 4% paraformaldehyde for 12 h. Tissues were dehydrated using 15% sucrose overnight at 4°C, and then 30% sucrose at 4°C for 2h. Tissue was contained in labeled embedding cassettes, and pre-cooled OCT was added to cover the entire tissue. Tissues were frozen quickly (10-20s) by placing the bottom of the box in contact with liquid nitrogen. Frozen tissue can be stored at −80°C. The tissues were cut into 8 μm slices. Sections were stained with H&E staining kit (C0105, Beyotime Biotechnology, China) and Masson trichrome staining kit (G1340, Solarbio, China) to assess histopathological changes in the lungs. Subsequently, all sections were observed under a microscope (DP73; Olympus, Tokyo, Japan)

### Western blot assay

After washing with cold PBS, 2 x 10^6^ cells were lysed in 1ml RIPA buffer (R0010-100, Solarbio) containing a protease inhibitor cocktail. 20 mg lung tissue was added to 100 μl of RIPA lysis buffer, and the tissue was first minced with tissue shears, then ground with zirconium beads (speed 12, time 4 min) and kept on ice for 30 mi. Total protein lysates were centrifuged at 15,000 × g for 15 minutes at 4°C. The supernatant was collected, and the total protein concentration was measured using the Bicinchoninic Acid (BCA) Protein Assay Kit (P1011, Beyotime Biotechnology, China). Total protein was diluted in SDS-PAGE loading buffer and denatured at 95°C for 10 minutes. Equal amounts of protein were loaded onto 10% SDS-PAGE gels and separated by electrophoresis. After electrophoresis, proteins were transferred to polyvinylidene difluoride (PVDF) membranes (Millipore, Burlington, MA). The membranes were blocked with 5% non-fat milk for 1 hour at room temperature, then incubated with primary antibodies against vimentin (1:1000), Fibronectin (1:1000), α-SMA (1:1000), Collagen I (1:2000), GAPDH (1:2000) overnight at 4°C. After washing three times with Tris-buffered saline containing 0.1% Tween 20, membranes were incubated with HRP-conjugated secondary antibodies for 1 hour at room temperature. ECL immunoblotting chemiluminescence solution was used for signal detection. An ultrasensitive multifunctional imager was used for exposure (BIO-RAD ChemiDoc Imaging System 733BR3967, USA).

### Real-time qPCR

Total RNA was extracted from A549 cells and MRC-5 cells using Trizol (Beyotime, Shanghai, China, R0016) and reverse transcribed using the Complementary DNA Reverse Transcription Kit (+gDNA wiper, R233-01, Vazyme, Nanjing, China). ChamQTM SYBR® qPCR Master Mix kit (High ROX Premixed, Q141-02, Vazyme Biotech Co. Ltd.) was used for amplification and quantitative fluorescence detection. The primers used are listed in Table 1. Gene expression values were calculated using the 2 ^−ΔΔCT^ method and normalized to GAPDH.

**Table 1.**
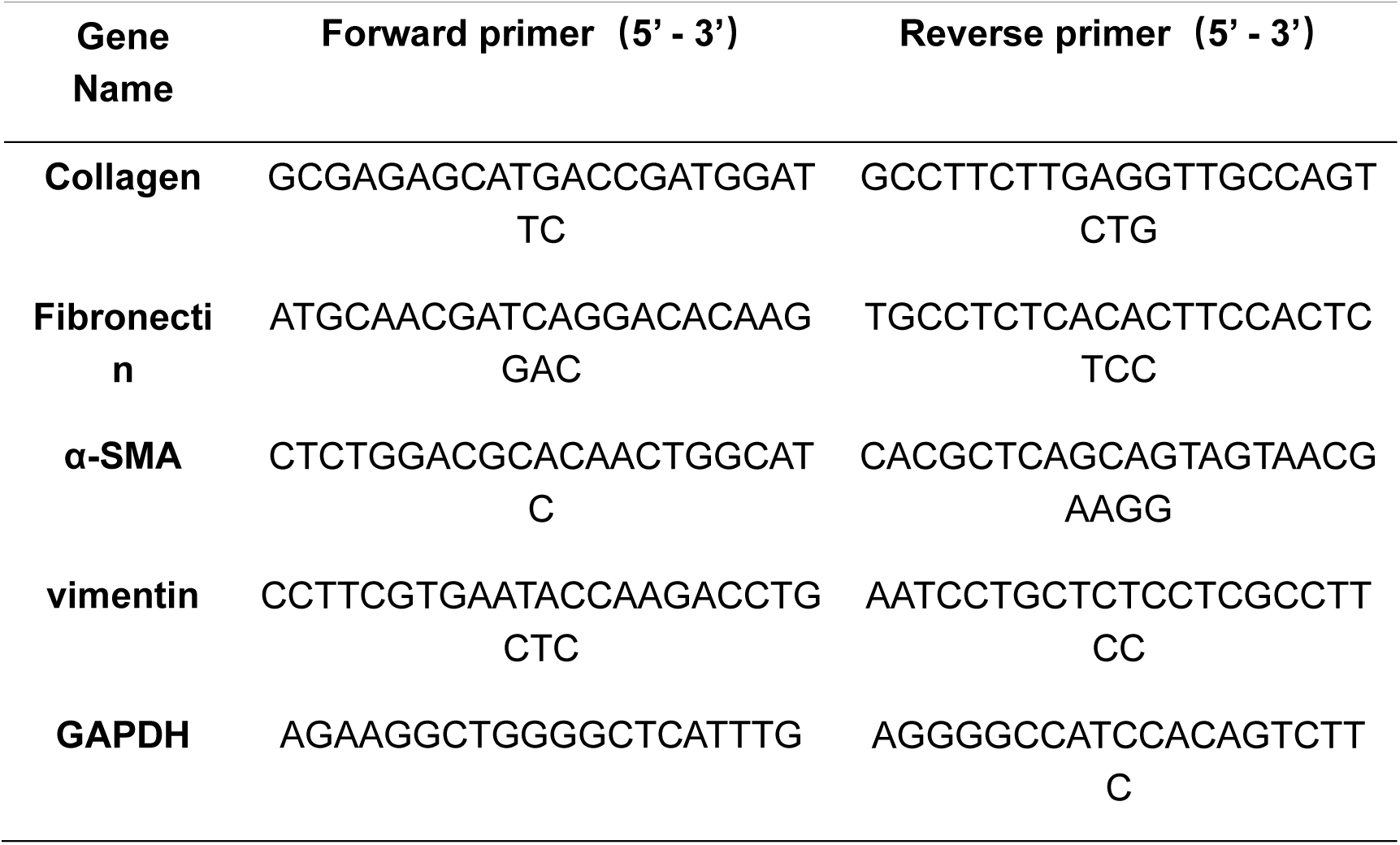
Primer sequences (Homo sapiens) for real-time qPCR analysis.

### Protein expression and purification

We developed a novel purification method for tag-free human vimentin based on its property of being denatured in a high-urea buffer, which increases its solubility. First, the no-tag vimentin plasmid was expressed in BL21*(DE3) Escherichia coli* cells and induced with 0.4 mM isopropyl β-D-thiogalactoside (IPTG, 0.4 M stock solution, 1:1000 dilution) at an OD600 of 0.7 at 37°C. Cells were pelleted by centrifugation at 5,000 × g for 30 minutes and resuspended in 50 ml of 10 mM Tris (pH 8.4) with 1 mM phenylmethylsulfonyl fluoride (PMSF, 100 mM stock solution, 1:100 dilution). Cells were lysed using a Low Temperature and Cell Disruptor (JN-Mini Pro, JNBIO, Guangzhou, China) at 1000–1200 mBar and 4°C for 3 cycles, until the lysate was no longer viscous. Vimentin, which is insoluble in the absence of urea, it was pelleted by centrifugation at 12,000 rpm for 60 minutes at 4°C. This step removed most of the soluble unwanted proteins. The pellet was then resuspended in a high-urea buffer (8 M urea, 10 mM Tris, pH 8.4) to solubilize vimentin. Another centrifugation (12,000 rpm, 30 minutes, 4°C) was performed to remove insoluble proteins and cell debris. The purification efficiency was evaluated using SDS-PAGE. In most cases, the purity of vimentin was sufficient for protein structural analysis. If the purity was not satisfactory, an additional purification step using a mixed-mode chromatography (MMC) column was employed. The MMC column was pre-equilibrated with 5 column volumes (CV) of ddH2O, followed by 10 CV of high-urea buffer (8 M urea, 10 mM Tris, pH 8.4). The filtered sample (0.45 μm filter) was then passed through the column twice to allow VIM to bind to the beads. The column was washed with high-urea buffer (8 M urea, 10 mM Tris, pH 8.4) to remove unbound proteins. vimentin was eluted using a gradient of ion-rich buffer (8 M urea, 1 mM EDTA, 5 mM MES, 1 M KCl, pH 6.0) from 0% to 100% on an AKTA PureMicro system (Cytiva). Fractions (0.5 ml) were collected in Eppendorf tubes, and UV280 absorbance was monitored. UV280 absorbance peaks were combined and concentrated. The purification efficiency was assessed using SDS-PAGE.

This was followed by the renaturation of vimentin : w with a 30-minute stepwise dialysis (6 M, 4 M, and 2 M urea in 5 mM Tris, 1 mM EDTA, 0.1 mM EGTA, 5 mM β-ME, pH 8.5) at room temperature to gradually reduce urea concentration. This was followed by an overnight dialysis in 0 M urea buffer (5 mM Tris, 1 mM EDTA, 0.1 mM EGTA, 5 mM β-ME, pH 8.5) at 4°C. Finally, vimentin exists as a tetramer after dialysis in a non-EDTA and non-EGTA buffer (5 mM Tris, 5 mM β-ME, pH 8.5) at room temperature for 1 hour. Proteins can be stored at −80℃.

### SPR analysis

The binding affinity of PD2 to vimentin was measured at 25°C using a Biacore X100 system (Cytiva). Briefly, compound solutions were prepared by serial dilution from stock solutions with running buffer. After completion of protein coupling to the CM5 chip, the ligand-fixed and reference channels were injected at 30 μl/min for 60 seconds (binding phase), followed by 120 seconds of dissociation monitoring. Data were analyzed using Biacore Insight Evaluation Software.

### Constructs

Virtual screening results indicate that PD2 forms eight hydrogen bonds with the target protein: its hydroxyl group forms seven hydrogen bonds with GLU138 in the A chain, GLU132 and GLU143 in the B chain, and GLU50 in the C chain; the oxygen atom on the tetrahydropyran ring forms one hydrogen bond with ARG54 in the C chain. To validate the interaction between PD2 and vimentin, we mutated all five key amino acid residues at the interface to Ala.

### Cellular Thermal Shift Assay (CETSA)

A549 cells were treated with 5 μM PD2 or DMSO for 24 hours. Cells were collected and resuspended separately in PBS containing PMSF. Aliquot the cell suspensions into 0.2 ml PCR tubes, with a volume of 100 μl per tube. Using a thermal cycler (Bio-Rad), heat the cells at 45°C–65.8°C for 3 minutes, then immediately freeze the cell suspensions rapidly in liquid nitrogen. Allow the tubes to stand at room temperature for 3 minutes. Perform two additional freeze-thaw cycles using the PCR machine and liquid nitrogen. Centrifuge the treated samples at 4°C at 12,000 × g for 20 minutes. Detect soluble proteins using Western blotting.

### In vitro assembly

Purified vimentin (1 mg/ml) was divided into two groups of 20 μl samples in 5 mM Tris, 5 mM β-ME, pH 8.5 buffer by gentle mixing. One group of Vimentin was incubated with a final concentration of 13.85 μM PD2, and the other (control) was treated with DMSO and incubated at RT for 30 min. After incubation, 20 μl of the Vimentin solution was transferred to an Eppendorf tube and preheated in a PCR thermocycler to 25°C. Filament formation was initiated by adding 20 μl of assembly buffer (40 mM Tris pH7.0,320 mM NaCl) and mixing rapidly. After 1 h, 40 μl of stop buffer (25 mM Tris pH7.5,50 mM NaCl,1 mM EDTA,0.1 mM EGTA,0.1 mM DTT) was added to each sample and mixed immediately.

### Defining thermostability of proteins by western blotting

To evaluate the thermal stability of purified recombinant vimentin, we performed a gradient heating western blot-based thermal stability assay as previously described with minor modifications^24^. Briefly, purified vimentin proteins or mutants (1 mg/ml) were assembled by incubation with PD2 or DMSO on ice for 1h. The assembled protein suspension was dispensed into 5 different 0.2 ml PCR tubes, each with a volume of 100 μl. Samples were heated at 45°C-65.8°C for 3 min using a thermal cycler (Bio-Rad), and then, immediately, the samples suspension was snap frozen in liquid nitrogen. Remove to room temperature for 3 minutes. Freeze-thaw the sample twice more using the PCR instrument and liquid nitrogen. The processed samples were centrifuged at 12,000 × g for 20 minutes at 4°C. Soluble proteins were detected by western blotting.

### Transmission electron microscopy of vimentin filaments

Negative staining was performed immediately after assembly to prepare EM samples. For negative staining, the glow-discharge carbon-coated copper grids were first rinsed with 4 μl of protein buffer (5 mM Tris, 5 mM β-ME, pH 8.5) and then with 4 μl of sample for 20 seconds. The samples were then fixed with 4 μl of 0.8% glutaraldehyde for 20 seconds. After fixation, the samples were stained twice with 4 μl uranyl acetate for 1 min 30 s each time. After the negatively stained samples were dried, the samples were examined using a Talos F200C transmission electron microscope and 12 representative images were collected from each group.

### Statistical analysis

GraphPad Prism v.8.0 was used to perform statistical analysis. Significance was determined by comparing two groups using the t-test analysis, and P < 0.05 was considered statistically significant. The data are represented as the mean ± SEM. The numbers of technical replicates or biological replicates for each group is indicated in the figure legends.

## Results

### PD2 is a potential ligand candidate of vimentin

To identify potential ligands targeting vimentin, we performed virtual screening over 70,430 compounds pooled from five libraries including four commercially available compound sets from MCE: a bioactive compound library (HY-L001V), a life chemical library (HY-L0107V), a protein mimetic library (HY-L0083V), a Chemspace scaffold-derived set (HY-L0093V) as well as an in-house tcmio ingredient library. Compounds were ranked by docking score, and the top candidates were selected for further analysis. Among them, PD2 — a triterpene saponin derived from *Platycodon grandiflorus* exhibited a notably favorable docking score of −10.634 kcal/mol (Figure 1A), indicating a robust thermodynamic binding potential toward the vimentin. To model the PD2–vimentin interaction, we constructed a vimentin tetramer by assembling two crystallographically resolved dimers according to the canonical coiled-coil packing mode. Molecular docking placed PD2 within a pocket at the intersubunit interface of the tetramer (Figure 1B).

**Figure 1.**
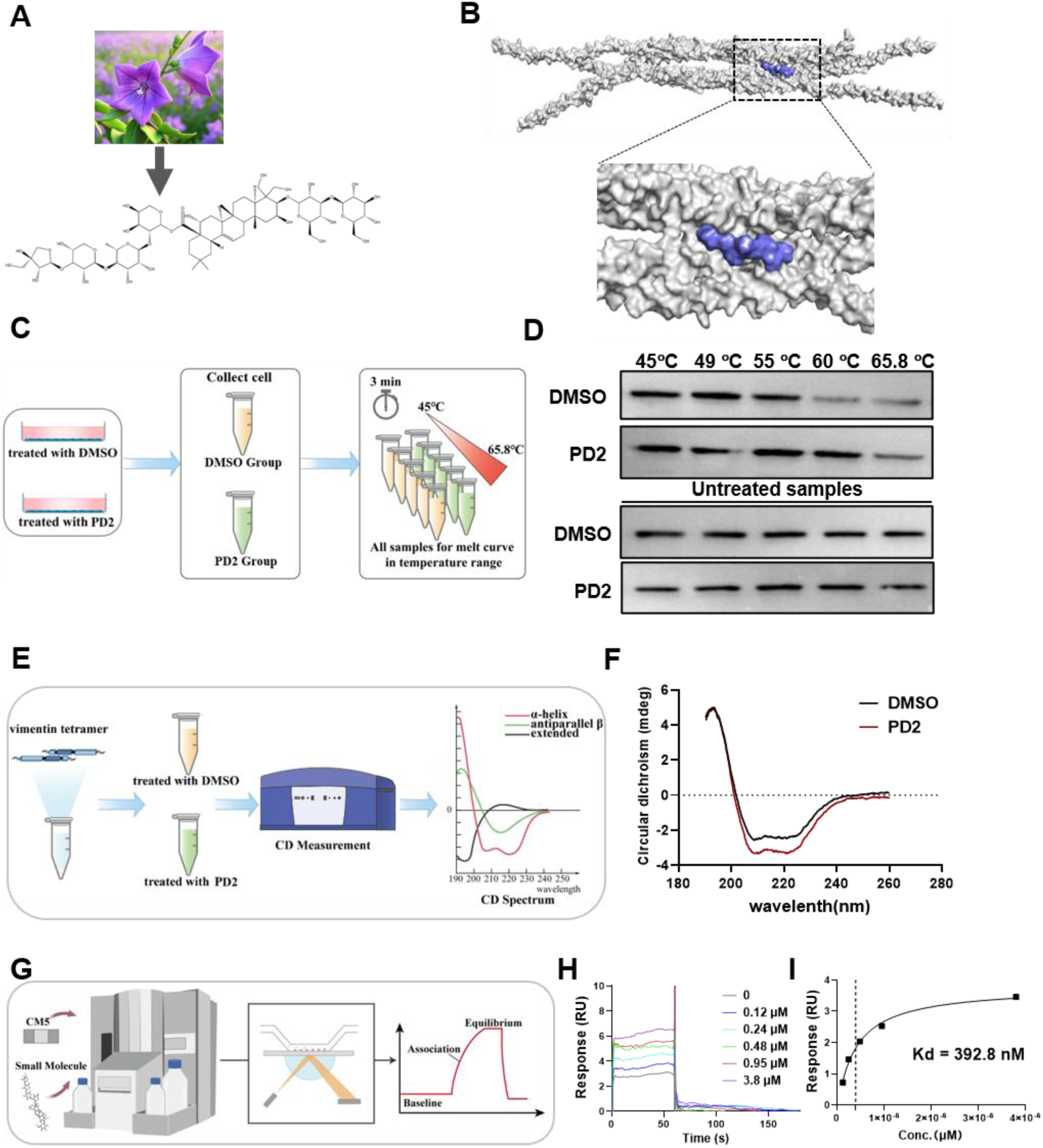
PD2 directly binds to vimentin proteins. **A** Chemical structure of PD2. **B** Molecular docking model of PD2-vimentin complex. The vimentin protein backbone is shown in gray and PD2 is depicted in light blue. **C/E/G** Schematic workflow of CETSA, CD, SPR assays used to detect the PD2-vimentin interaction, respectively. **D** CETSA results showing that PD2 stabilized the vimentin compared with DMSO control group in A549 cells, as detected by anti-vimentin. **F** CD analysis demonstrating that PD2 induced changes in the secondary structure content of vimentin. **H-I** SPR sensorgram and binding affinity determination, yielding a measured Kd of 392.8 nM between PD2 and vimentin.

To verify this interaction, we adopted several experimental assays. First, a cell-based thermal shift assay (CETSA) was performed in A549 lung cancer cells (Figure 1C). Compared with the DMSO-treated control, PD2 treatment markedly enhanced the thermal stability of vimentin (Figure 1D), indicating that PD2 stabilizes vimentin in a cellular context. We applied circular dichroism (CD) spectroscopy to monitor conformational changes in protein secondary structure (Figure 1E). CD analysis further showed that PD2 induces an increase in the α-helix content of vimentin, which indicates this shift in secondary structure is induced by PD2-vimentin interaction (Figure 1F). Finally, quantitative determination of the binding affinity between vimentin and PD2 using surface plasmon resonance (SPR) (Figure 1G-I). The result showed a Kd value of 392.8 nM (Figure 1H, I). Taken together, these results demonstrate that PD2 directly binds to vimentin with high affinity and alters its secondary structure both in vitro.

### PD2 attenuates vimentin expression in lung epithelial cells

To establish a suitable concentration range for functional studies, we first evaluated the cytotoxic profile of PD2 in A549 lung epithelial cells. Across a gradient of concentrations, cell viability remained fully preserved at 6 μM PD2 (Figure 2A), and a half-maximal inhibitory concentration (IC₅₀) was calculated to be 16 μM (Figure 2B). Based on these results, subsequent cellular experiments were conducted using a working concentration of 5 μM PD2.

**Figure 2.**
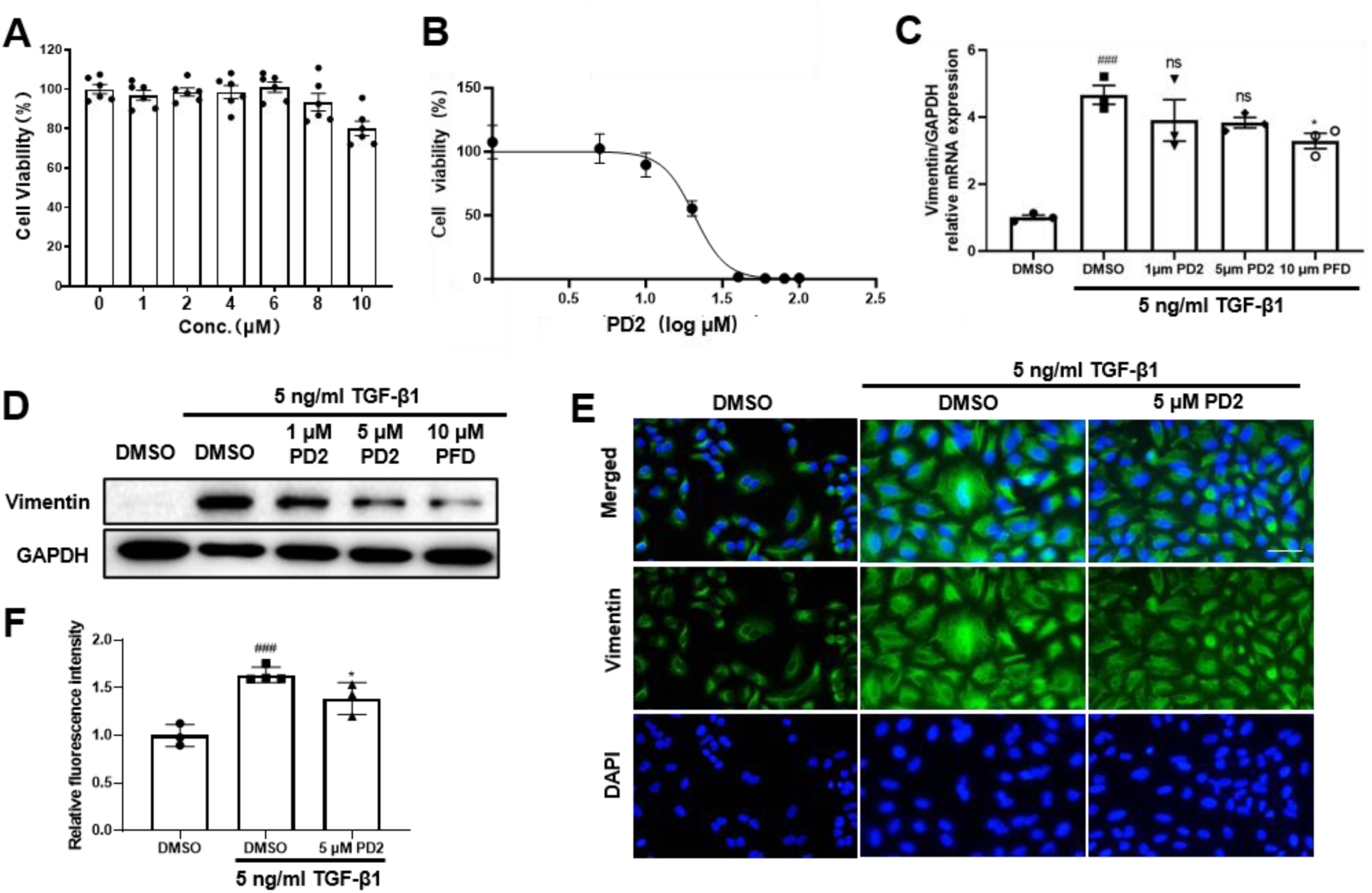
PD2 inhibits vimentin expression in TGF-β1-induced A549 cells. **A** Cell viability of A549 cells following PD2 treatment at the indicated concentrations, as determined by CCK-8 assay. **B** Determination of the half-maximal inhibitory concentration (IC50) of PD2 in A549 cells. **C** qRT-PCR analysis of vimentin mRNA levels in TGF-β1-induced A549 cells following PD2 or PFD intervention. Data are expressed as mean ± SEM (n = 3). ^###^P < 0.001 versus the DMSO control group. *P < 0.05 versus the DMSO with TGF-β1 group. **D** Western blot analysis of vimentin protein levels in TGF-β1-induced A549 cells 48 hours after PD2 or PFD treatment. **E** Representative immunofluorescence staining to assess the fluorescence intensity of intracellular vimentin in TGF-β1-induced A549 cells following PD2 intervention. Scale bar: 50 µm. **F** Quantitative analysis of vimentin immunofluorescence intensity in (E). Data are expressed as mean ± SEM (n = 3-4). ^###^ P < 0.001 compared with the DMSO control group. *P < 0.05 compared with the DMSO + TGF-β1 group.

To examine the effect of PD2 on vimentin under fibrotic conditions, we stimulated A549 cells with TGF-β1, a well-characterized inducer of epithelial–mesenchymal transition (EMT)^15^ that upregulates vimentin expression during this process. We first quantified VIM mRNA levels by qPCR. Unlike pirfenidone (PFD), a clinically approved antifibrotic agent, PD2 treatment did not cause any appreciable change in transcript level (Figure 2C). In contrast, Western blot analysis revealed that PD2 reduced vimentin protein to a level comparable to that of PFD (Figure 2D). This finding was independently confirmed by immunofluorescence staining (Figure 2E). Taken together, these results suggest that PD2 may influence overall vimentin levels by affecting its post-transcriptional regulation, which is mechanistically distinct from that of PFD.

### PD2 blocks FMT in fibroblast

Having established that PD2 downregulates vimentin at the protein level, we next asked whether this reduction suppresses the FMT, a central pathological process in fibrosis driven in part by vimentin upregulation^25^. To this end, we employed TGF-β1-stimulated MRC-5 human fetal lung fibroblasts as an in vitro FMT model for evaluating the anti-fibrotic effects of PD2.

We evaluated the anti-fibrotic effects of PD2 using multiple FMT-associated strategy. Upon TGF-β1 stimulation, quiescent fibroblasts undergo phenotypic conversion into myofibroblasts, characterized by increased expression of α-SMA, fibronectin, and collagen, as well as enhanced migratory capacity and extracellular matrix secretion. Scratch wound healing assays demonstrated that PD2 treatment significantly suppressed TGF-β1-induced cell migration (Figure 3A). Consistent with this observation, both qPCR and western blot analysis revealed that PD2 effectively downregulated the expression of key FMT-related genes, including ACTA2 (encoding α-SMA) and FN1 (fibronectin) (Figure 3B, C, D). Furthermore, PD2 markedly reduced intracellular hydroxyproline content (Figure 3E), directly demonstrating its ability to attenuate collagen deposition—a key determinant of tissue stiffness and lung function decline in pulmonary fibrosis.

**Figure 3.**
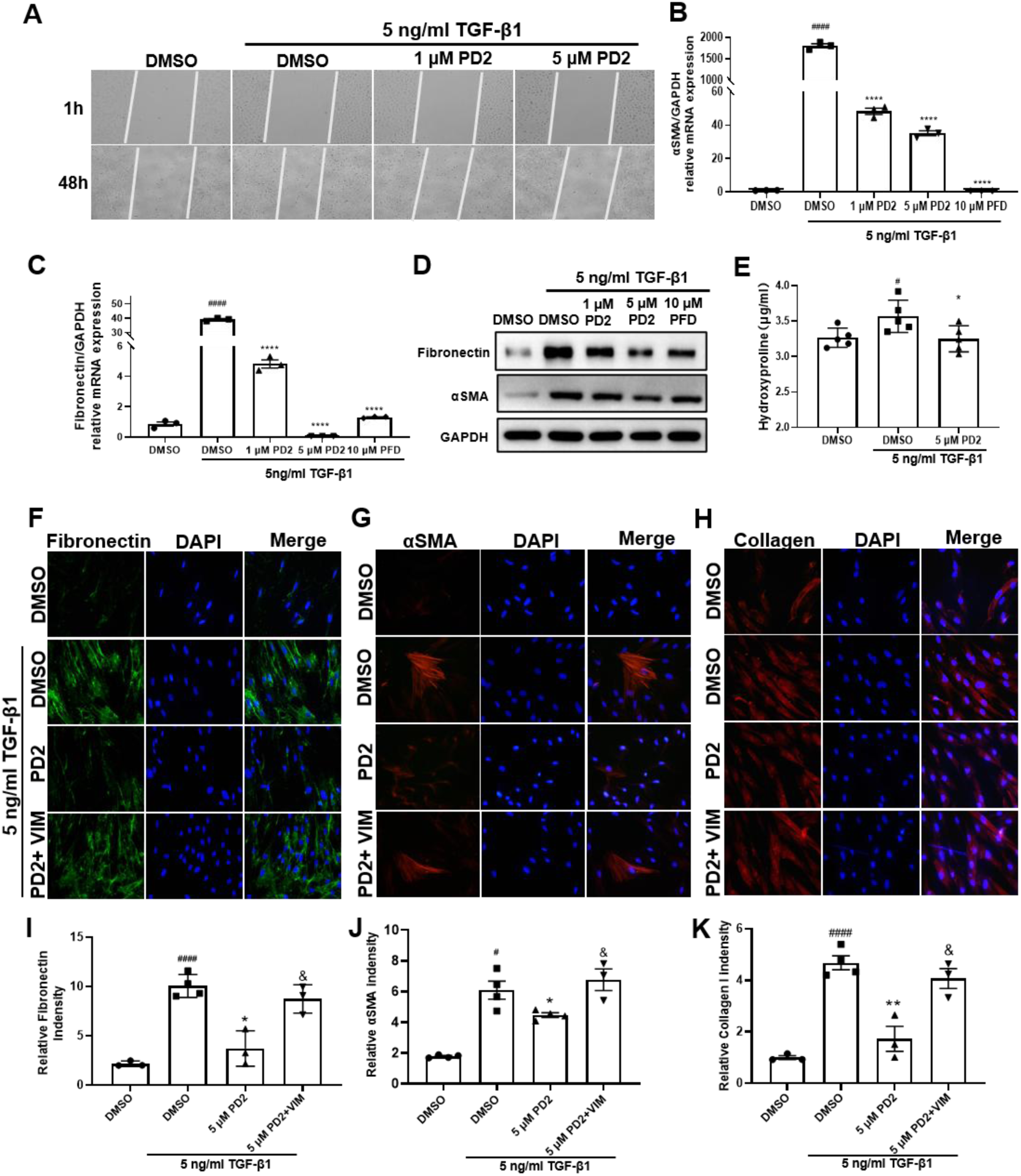
PD2 attenuates FMT in TGF-β1-stimulated MRC-5 cells. **A** Scratch wound healing assay showing that PD2 treatment inhibited TGF-β1-induced migration of MRC-5 cells. **B-C** qPCR analysis showing that PD2 treatment reduced the mRNA levels of fibrosis-related genes in TGF-β1 stimulated MRC-5 cells. **D** Western blot analysis showing that PD2 treatment decreased the protein expression levels of fibrosis markers under the same conditions as in C. **E** Measurement of intracellular hydroxyproline content reflecting a marked reduction of collagen deposition following PD2 treatment in TGF-β1-stimulated MRC-5 cells. **F–H** Immunofluorescence staining showing decreased intracellular fibronectin, α-SMA, and collagen upon PD2 treatment. Scale bar: 50 µm. **I–K** Quantification of immunofluorescence intensity in F-H demonstrating that exogenous vimentin supplementation partially attenuated the anti-fibrotic effects of PD2. ^#^P < 0.05, ^####^P < 0.0001, versus the DMSO control group. *P < 0.05, **P < 0.01, ****P < 0.0001, versus the DMSO + TGF-β1 group. ^&^ P < 0.05, versus the 5 μM PD2 group.

If PD2 exerts its anti-fibrotic effects primarily through vimentin downregulation, then restoring vimentin levels should at least partially rescue the FMT phenotype. To test this prediction, we performed a vimentin rescue experiment by supplemented PD2-treated fibroblasts with exogenous vimentin and assessed the FMT marker expression levels by immunofluorescence staining (Figure 3F-H) and quantitative analysis (Figure 3I-K). Exogenous vimentin supplementation partially reversed the PD2-mediated suppression of expression (Figure 3F-K). Collectively, these results demonstrate that PD2 blocks FMT in lung fibroblasts by reducing vimentin levels.

### PD2 ameliorates bleomycin-induced pulmonary fibrosis in vivo

To evaluate the anti-fibrotic efficacy of PD2 in vivo, as described earlier^26,27^, a bleomycin-induced pulmonary fibrosis model in mice was established following the experimental schedule outlined in Figure 4A. PD2 treatment improved the survival rate of fibrotic mice (Figure 4B) and attenuated the progressive body weight loss associated with bleomycin-induced lung injury (Figure 4C). Hematoxylin and eosin (H&E) staining revealed that PD2 treatment markedly reduced inflammatory cell infiltration and alveolar structural disruption, while Masson’s trichrome staining demonstrated a substantial decrease in collagen deposition (Figure 4D). Semi-quantitative scoring of lung tissue sections indicated that following PD2 administration, both the Ashcroft score (reflecting the severity of fibrosis) and the H&E score (reflecting inflammatory damage) were significantly reduced (Figure 4E-F). Consistent with these findings, the hydroxyproline content in lung tissue, a well-established biochemical index of total collagen burden, was significantly reduced in PD2-treated mice compared with vehicle-treated controls (Figure 4K). Consistent with this reduction in extracellular matrix deposition, western blot analysis confirmed the downregulation of key fibrotic markers, including α-SMA and fibronectin (Figures 4G–J). Taken together, these results demonstrate that PD2 effectively alleviates bleomycin-induced pulmonary fibrosis in mice, supporting its anti-fibrotic potential in vivo.

**Figure 4.**
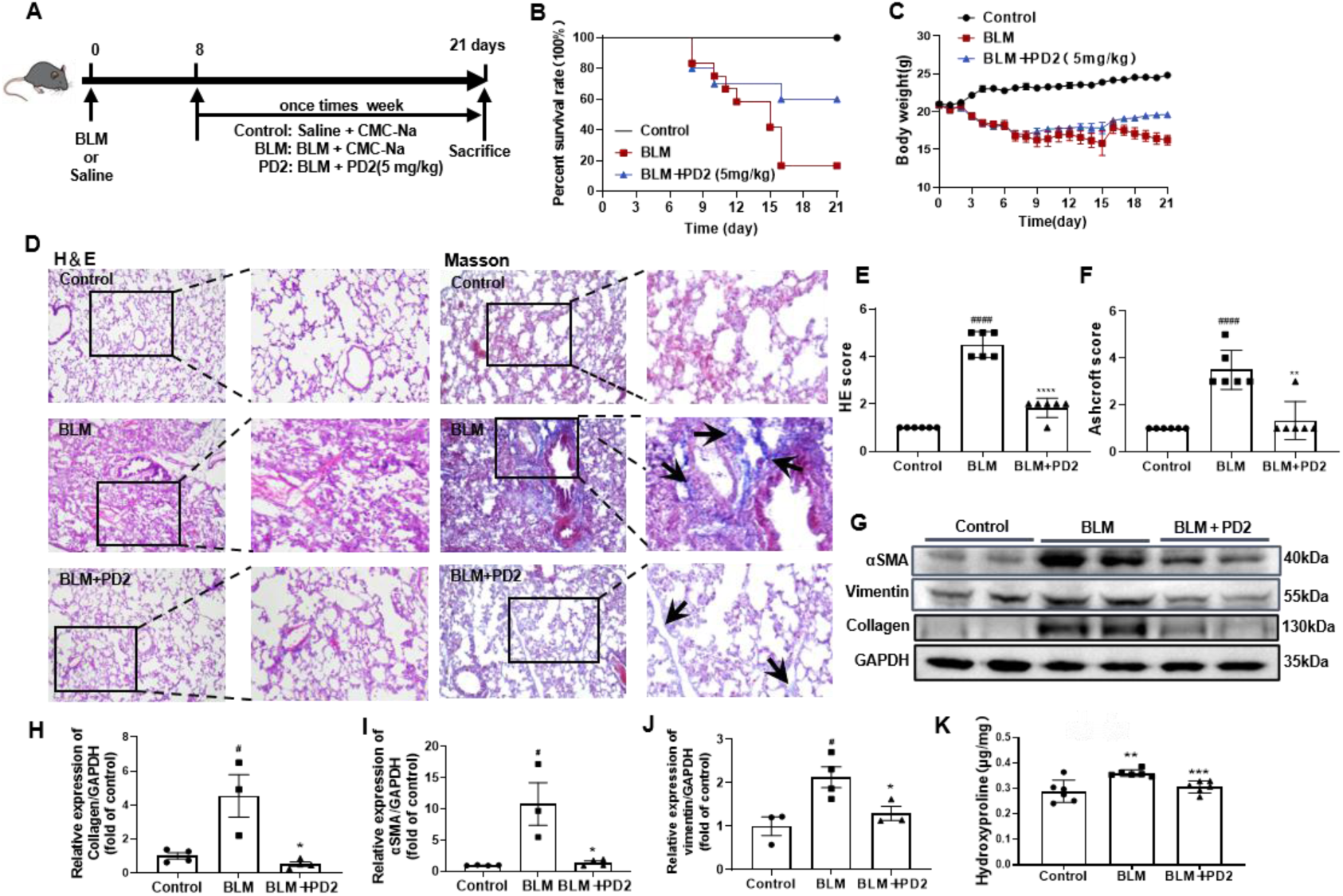
PD2 ameliorates bleomycin-induced pulmonary fibrosis in mice. **A** Flowchart of the bleomycin-induced pulmonary fibrosis animal model. C57BL/6J mice were treated with bleomycin: on Day 0, they received a single intratracheal infusion of bleomycin (5 mg/kg); starting on Day 8, they received weekly treatments with CMC-Na (control group) or PD2; and on Day 21, they were euthanized (n = 6 mice per group). **B** Mouse survival rate showing that PD2 treatment improved the survival rate of bleomycin-treated mice. **C** Body weight measurements showing that PD2 treatment attenuated bleomycin-induced weight loss. **D** Representative images of lung tissue sections stained with hematoxylin and eosin (H&E) and Masson’s trichrome, showing that PD2 treatment reduced inflammatory infiltration, alveolar structural disruption, and collagen deposition. **E-F** Quantitative scoring of lung tissue sections, showing significantly reduced H&E score and Ashcroft score following PD2 administration (n = 6). **G** Western blot analysis showing decreased protein levels of collagen, α-SMA, and vimentin in lung tissue from PD2-treated mice. **H-J** Quantification of Western blot band intensities in G by ImageJ (n=3-6). Data are presented as mean ± standard deviation. **K** Measurement of hydroxyproline content in lung tissue, showing a marked reduction following PD2 treatment. ^#^P < 0.05, ^##^P < 0.01, ^####^P < 0.0001 versus the control group; *P < 0.05, **P < 0.01, ***P < 0.001,****P < 0.0001 versus the BLM group.

### PD2 induces aberrant aggregation of vimentin

The discordance between reduced vimentin protein levels and unchanged *Vimentin* mRNA abundance (Figure 2) suggested that PD2 acts through a post-transcriptional mechanism. Given that PD2 directly binds to vimentin and increases its α-helical content (Figure 1), we reasoned that this binding event might perturb the normal assembly dynamics of vimentin. Vimentin function depends on its ability to undergo regulated cycles of assembly into insoluble intermediate filaments and disassembly into soluble tetramers. A ligand-induced conformational shift could, in principle, disrupt this equilibrium, diverting the protein toward an aberrantly aggregated state^8^. To test this hypothesis, we examined the solubility state of vimentin following PD2 treatment. Immunofluorescence staining revealed that levels of insoluble vimentin were significantly elevated 48 h after PD2 treatment (Figure 5A). Western blot analysis further confirmed the accumulation of insoluble vimentin in PD2-treated cells (Figure 5B). These results indicate that PD2 binding promotes the conversion of vimentin from a soluble, functional pool into an insoluble, aggregated state in cells.

**Figure 5.**
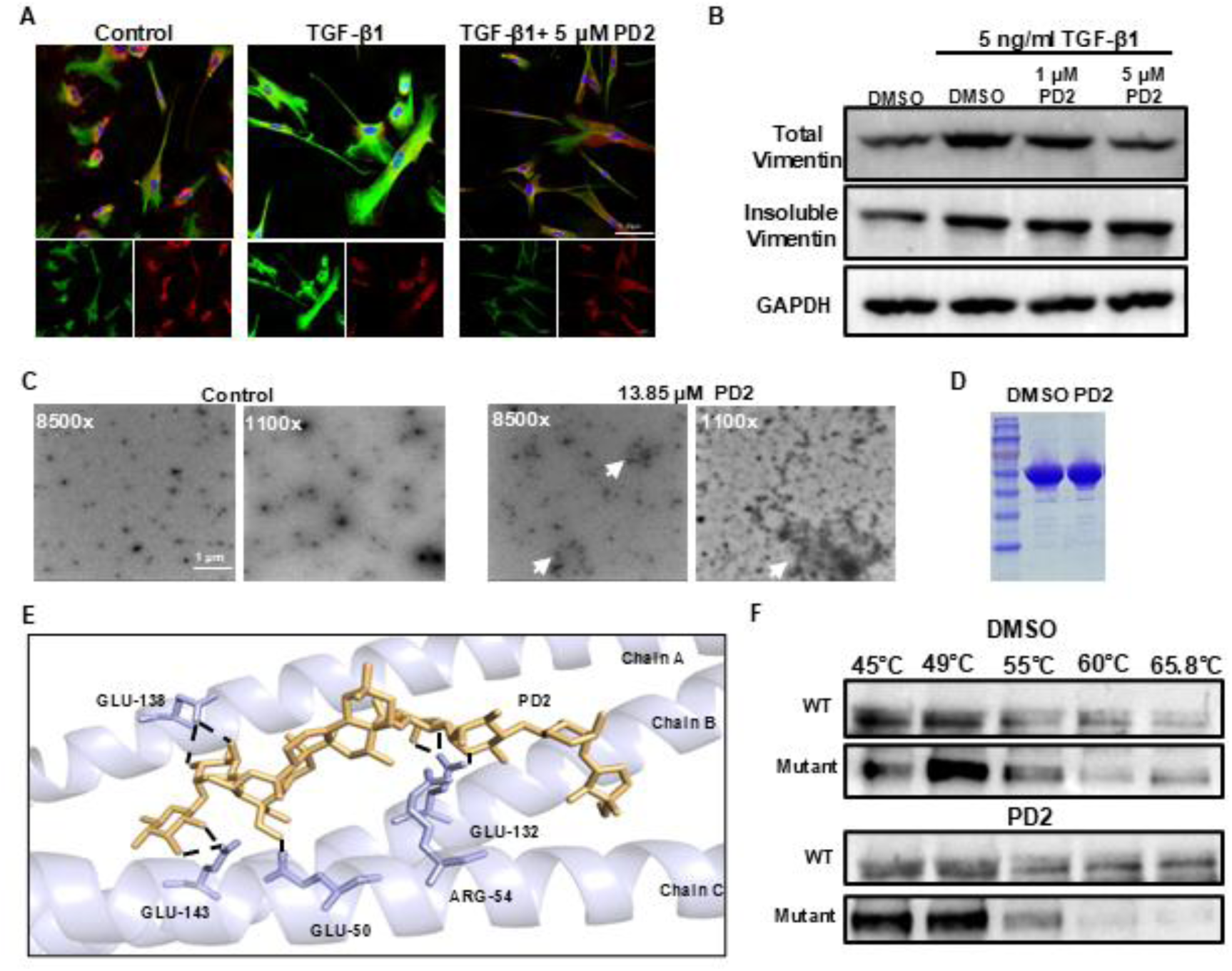
PD2 targets vimentin and promotes its aggregation. **A** Immunofluorescence staining results showing a marked accumulation of insoluble vimentin increase 48 h after PD2 treatment (scale bar: 20 µm). **B** Western blot analysis further confirms the accumulation of insoluble vimentin in PD2-treated cells. **C** In vitro vimentin assembly assay demonstrating that PD2 directly induces vimentin aggregation relative to the DMSO vehicle control. **D** SDS-PAGE analysis confirming equivalent protein loading and purity across conditions shown in C. **E** Molecular docking simulations showing the predicted binding mode of PD2 to vimentin and the key amino acid residues potentially involved in the vimentin-PD2 interaction were shown. **F** Thermal stability assays showed that mutations at potential interaction sites in vimentin eliminate its interaction with PD2.

To determine whether this aggregation is a direct consequence of PD2 binding or an indirect cellular response, we performed a cell-free vimentin assembly assay using purified vimentin protein. PD2, but not the vehicle control, directly triggered vimentin aggregation in vitro (Figure 5C), demonstrating that PD2 is sufficient to induce this conformational transition without the requirement of additional cellular factors.

We next sought to identify the structural basis of the PD2–vimentin interaction. Molecular docking simulations revealed that PD2 occupies a pocket formed by residues located at the interface of vimentin coiled-coil domains, where it is stabilized through multiple intermolecular interactions. Specifically, the hydroxyl groups of PD2 form hydrogen bonds with the side chains of Glu138 in chain A, Glu132 and Glu143 in chain B, as well as Glu50 and Arg54 in chain C (Figure 5E). To test whether these residues are required for PD2 binding, we generated a vimentin mutant in which all five candidate residues were substituted with alanine. CETSA experiments revealed that these site-directed mutations completely abolished the ligand-induced thermal stabilization of vimentin by PD2 (Figure 5F), indicating that these specific residues are required for the PD2–vimentin interaction. Collectively, these findings suggest that PD2 directly binds to vimentin, inducing a conformational change that redirects vimentin from its native state into an insoluble, aberrantly aggregated state, which functionally inactivates this pro-fibrotic protein(Figure 6). This aggregation-dependent inactivation, rather than degradation-dependent clearance, represents a distinct mechanistic strategy for targeting vimentin in fibrosis.

**Figure 6:**
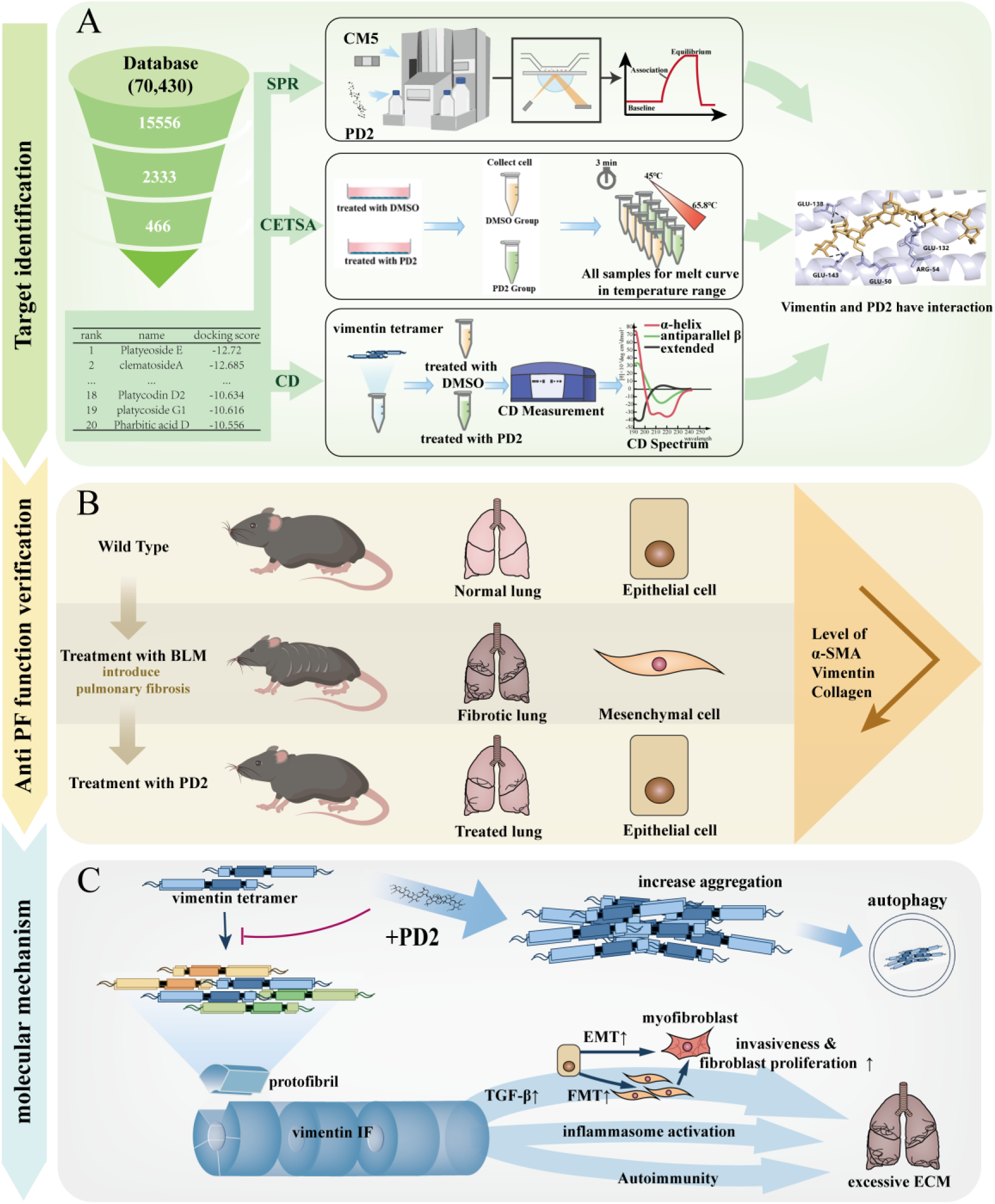
Schematic summary of this study. **A** Target identification. PD2 was identified from virtual screening of the TCMIO database and its direct interaction with vimentin was validated by SPR, CETSA, and CD assays. **B** Anti-pulmonary fibrosis function. PD2 treatment ameliorated bleomycin-induced pulmonary fibrosis in mice, as evidenced by restored lung architecture and reduced levels of α-SMA, vimentin, and collagen. **C** Molecular mechanism. PD2 promotes vimentin tetramer aggregation, thereby inhibiting its assembly into intermediate filaments and suppressing downstream pathological processes including EMT, FMT, and excessive ECM deposition to attenuate pulmonary fibrosis.

## Discussion

PF is a chronic and irreversible disease that progressively leads to respiratory failure and death^28^. The abnormal expression and activation of vimentin play a crucial role in PF pathogenesis. Notably, vimentin intermediate filaments are significantly upregulated in myofibroblasts within fibrotic areas of patients with idiopathic pulmonary fibrosis (IPF) ^29^. As a core member of the intermediate filament protein family, vimentin is critically involved in mechanical stress transmission, amplification of inflammatory signals, and cross-talk with the TGF-β pathway^20^, making it an attractive therapeutic target for PF. Although targeting vimentin has emerged as a potential anti-fibrotic strategy, no vimentin-specific inhibitor has been approved for clinical use^9^. Thus, developing novel agents that specifically target and modulate vimentin function is an urgent and promising research direction.

Here, through virtual screening, we identified PD2 as a top-ranking compound binding to vimentin. While previous studies suggested the empirical anti-fibrotic clinical potential of PD2 via TGF-β/Smad and NF-κB pathways^30^, its direct intra-cellular receptor remained elusive. Our biophysical evidence (CETSA, SPR, and CD) successfully bridges this gap by positioning vimentin as a high-affinity functional target of PD2. Notably, qRT-PCR analysis revealed that PD2 had virtually no effect on *vimentin* mRNA expression, suggesting that PD2 may ultimately regulate vimentin protein levels by influencing post-transcriptional processes. We then explored the anti-fibrotic mechanism of PD2 by detecting FMT markers such as fibronectin and α-SMA. We found that PD2 effectively inhibits TGF-β1-induced fibrosis, thereby blocking the conversion of fibroblasts into myofibroblasts and reducing collagen deposition. In a bleomycin-induced murine model of PF, PD2 treatment exhibited significant anti-fibrotic effects: it improved survival, reduced weight loss, decreased collagen accumulation, and downregulated key fibrotic markers. These in vivo results strongly support vimentin-targeting as a viable therapeutic strategy.

To elucidate the mechanism underlying PD2-mediated vimentin downregulation, we investigated the fate of vimentin following PD2 binding. Immunofluorescence, biochemical sedimentation, and in vitro assembly assays revealed that PD2 binds specific residues of vimentin, inducing the formation of insoluble aggregates and disrupting vimentin function. Our study identifies a novel mechanism whereby PD2 promotes the formation of insoluble filamentous vimentin aggregates. This mechanism differs from known vimentin-targeting inhibitors. Withaferin A causes vimentin degradation via covalent modification^17^, while FiVe1 inhibits vimentin polymerization^31^. In contrast, PD2 does not eliminate vimentin through canonical degradation pathways but instead traps it in an inactive aggregated state—a mechanism shared with the natural compound ATD, which also stabilizes vimentin multimers and disrupts function in PF models^20^. This supports the therapeutic potential of pharmacologically inducing vimentin aggregation.

However, it is important to note that protein aggregation is often associated with cytotoxicity and neurodegenerative diseases. We speculate that intermittent dosing or concentrations below the IC50 of PD2 may confine vimentin aggregation within a controllable and safe window, specifically disrupting pathological vimentin function without significant toxicity^32^. Consistent with this view, we observed no cytotoxicity at the working concentration of 5 μM in cell-based assays (Figure 2A). Nevertheless, this hypothesis requires further validation through in vivo toxicological studies. Molecular docking and mutagenesis study further indicated that PD2 likely binds vimentin to interfere with its normal function. Future structural studies will be needed to define the precise binding mode at atomic resolution.

We propose the following mechanistic model: PD2 directly binds vimentin, disrupts its normal assembly dynamics, and induces aberrant aggregation. This dysfunctional state subsequently interferes with the core fibrotic driver—TGF-β1 signaling cascade—ultimately inhibiting FMT and extracellular matrix synthesis. This model is consistent with previous reports highlighting the functional interplay between vimentin and the TGF-β1 signaling pathway^33^. Future studies should prioritize systematic in vivo toxicity evaluations, including maximum tolerated dose, histopathological assessment of major organs, and evaluation of potential neurotoxicity. Furthermore, elucidating the structural basis of PD2-vimentin interaction at high resolution and evaluating the efficacy of PD2 in other preclinical models of fibrosis are essential next steps toward clinical translation.

## Conclusion

In summary, our study reveals a novel anti-fibrotic mechanism of PD2: it binds to vimentin and induces of insoluble aggregates, thereby inactivating its pro-fibrotic function. This finding not only underscores the therapeutic potential of vimentin-targeting strategies but also establishes PD2 as a promising lead compound and chemical probe for further investigating the role of vimentin in fibrosis.

## Author statement

X. L.:Writing-original draft, visualization, validation, methodology, investigation, formal analysis, data curation. C.Y.: Visualization, investigation, formal analysis. X.M., Y.K. and investigation, formal analysis. W.W.: Resources. S.S.: Writing—review and editing, supervision, project administration, funding acquisition, conceptualization. All authors have read, agreed to the published version of the manuscript, and assured the accuracy of investigation.

## Funding

This research was funded by the Natural Science Foundation of Shandong Province, China (ZR2022QC213), National Natural Science Foundation of China (32301014).

## Institutional Review Board Statement

The animal study protocol was approved by the Ethics Committee of School of Basic Medical Sciences, Shandong University (protocol code: ECSBMSSDU2023-2-114, date of approval: 24 Feb 2023).

## Data Availability Statement

The original contributions presented in this study are included in the article. Further inquiries may be directed to the corresponding authors.

## Declaration if Competing Interest

The authors declare that they have no competing interests.

## Clinical trial number

Not applicable.

## Consent to Publish declaration

Not applicable.

## Consent to Participate declaration

Not applicable.

## Notes

### Competing Interest Statement

The authors have declared no competing interest.

